# Dissociation of Molecular and Behavioral Neuroadaptations Following Acute GRK2/3 Inhibition in Amphetamine-Treated Rats

**DOI:** 10.64898/2026.04.14.718549

**Authors:** Catalina Starocelsky, María Estela Andrés, Rafael Ignacio Gatica

**Author notes:** current address (Rafael Ignacio Gatica).

## Abstract

**Background:** Individual vulnerability to addiction is driven by neuroadaptations within dopaminergic circuits. G protein-coupled receptor kinases (GRKs), specifically GRK2 and GRK3 regulate D2 receptor (D2R) signaling and trafficking, but their role in amphetamine (AMPH)-induced locomotor sensitization remains unclear. This study aimed to determine whether GRK2/3 inhibition alters locomotor sensitization, and its associated molecular correlates across striatal regions.

**Methods:** Adult rats (n = 39) were assigned to saline, acute AMPH, or repeated AMPH groups and received intraperitoneal administration of vehicle or the GRK2/3 inhibitor Cmpd101 (1.0 mg/kg intraperitoneally, i.p.). Locomotor activity was assessed under basal and injection conditions to evaluate sensitization. Protein levels of D2R, GRK2, and GRK5 were quantified across striatal regions using Western blot and analyzed with linear mixed models.

**Results:** Cmpd101 did not significantly affect the acute hyperlocomotor effects of AMPH or the expression of AMPH sensitization. At the molecular level, Cmpd101 had no effect on D2R levels and produced selective, region-dependent changes in GRK2 and GRK5. Notably, GRK2/3 inhibition altered the relationship between protein expression and the degree of locomotor sensitization in a region-specific manner, rather than inducing consistent changes in absolute protein levels.

**Conclusions:** GRK2/3 inhibition by Cmpd101 produces region-specific molecular effects and reshapes protein-behavior relationships without significantly altering locomotor sensitization. These findings support a model in which GRKs act as context-dependent modulators of dopaminergic signaling rather than direct drivers of behavioral output.

## 1 Introduction

Drug addiction is a chronic brain disease that entails severe health and socioeconomic consequences worldwide (Degenhardt & Hall, 2012). It is characterized by compulsive drug use despite adverse consequences (Koob & Volkow, 2016). Individual vulnerability to addiction is a complex phenomenon influenced by behavioral, environmental, cellular, molecular, and genetic factors (Anderson & Hearing, 2019; Egervari et al., 2018; Hamer, 2002). Importantly, drug use does not necessarily lead to addiction, and not all individuals who consume drugs become addicted. It is estimated that approximately 15-17% of individuals who consume addictive substances transition from recreational use to chronic drug consumption (Anthony et al., 1994; Lopez-Quintero et al., 2011). This observation suggests that certain individuals are more vulnerable to the reinforcing effects of drugs of abuse (Egervari et al., 2018; Piazza & Deroche-Gamonet, 2013). Such individual differences in vulnerability have also been consistently observed in animal models of addiction (Anker et al., 2009; Belin et al., 2008; Belin & Deroche-Gamonet, 2012; Bush & Vaccarino, 2007; Deroche-Gamonet et al., 2004; Kuhn et al., 2022; Marinelli & White, 2000; Piazza et al., 1989; Piazza et al., 2000; Swain et al., 2021; Volkow & Morales, 2015).

Repeated psychostimulant exposure can induce locomotor sensitization, an enduring enhancement of drug-induced hyperlocomotion, widely used as a preclinical model to study persistent adaptations linked to vulnerability and relapse (Pierce & Kalivas, 1997; Steketee & Kalivas, 2011). Amphetamine (AMPH) elevates extracellular dopamine (DA), primarily by reversing dopamine transporter (DAT) function and blocking the vesicular monoamine transporter 2 (VMAT2) (Robertson et al., 2009; Sulzer, 2011; Teng et al., 1998; Volkow et al., 2011), and expression of locomotor sensitization to AMPH is accompanied by region-specific alterations in dopaminergic transmission (Di Chiara & Imperato, 1988; Gatica et al., 2020; Scholl et al., 2009; Sullivan et al., 1998). The resulting increase in extracellular DA enhances activation of dopaminergic receptors, which belong to the G protein-coupled receptor (GPCR) family and mediate intracellular signaling cascades critical for synaptic plasticity and behavioral adaptation (Thompson et al., 2010). Within this framework, dopamine D2 receptors (D2R), expressed at both pre- and postsynaptic sites, are key modulators of DA signaling and have been implicated in addiction-related phenotypes (Ashok et al., 2017; Bello et al., 2011; Schmitz et al., 2001).

Upon agonist activation, G protein-coupled receptor kinases (GRKs) orchestrate the processes of GPCR desensitization, internalization, and recycling (Gurevich & Gurevich, 2020). In particular, GRK2 and GRK3 are central regulators of D2R signaling and trafficking (Kim et al., 2001; Mann et al., 2021; Namkung et al., 2009a; Namkung et al., 2009b; Pack et al., 2018). GRK2 is broadly expressed in reward-related brain regions, including colocalization with striatal medium spiny neurons and dopaminergic neurons, whereas GRK3 expression appears more restricted, suggesting distinct contributions to D2R regulation (Erdtmann-Vourlioti et al., 2001). Psychostimulants exposure has been associated with changes in GRK expression in reward circuits, yet reported behavioral effects of GRK2/3 manipulations are inconclusive (Daigle et al., 2014; Gainetdinov et al., 2004; Sellgren et al., 2021). Notably, the selective GRK2/3 inhibitor Cmpd101 (Thal et al., 2011) reduces GRK2-D2R interactions and blocks dopamine-induced D2R internalization (Sánchez-Soto et al., 2023), highlighting GRK2/3 as potential modulators of DA-dependent behaviors. However, whether GRK2/3 inhibition alters basal and AMPH-induced locomotor responses, including sensitization-related phenotypes, remains unclear. Therefore, the present study examined the effects of GRK2/3 inhibition with Cmpd101 on basal and AMPH-induced locomotor activity in rats. Our data showed that while Cmpd101 modulates protein levels and protein-behavior relationships in a region-specific manner, these effects are not translated into robust changes in locomotor behavior after acute or repeated AMPH.

## 2 Material and methods

### 2.1. Animals

Rats were obtained from the Animal Care Facility of the Biological Sciences at the Pontificia Universidad Católica de Chile. They were housed in a colony room in groups of two per cage and kept at room temperature between 20 and 24 °C on a 12 h light/dark cycle (lights on at 9 AM), with access to food and water ad libitum. All protocols were approved by the local bioethics committees (Project #220420014, Pontificia Universidad Católica de Chile). All procedures were in strict accordance with the guidelines published in the “NIH Guide for the Care and Use of Laboratory Animal” (8th Edition) and the principles presented in the “Guidelines for the Use of Animals in Neuroscience Research” by the Society for Neuroscience. Rats were handled for 4 days before starting the experiments. A total of 39 rats were used in this study.

### 2.2. Drugs

Animals received intraperitoneal (i.p.) administrations of the following solutions. Saline-treated groups were injected with 0.9% NaCl (Laboratorio Sanderson S.A., Santiago, Chile). AMPH was administered as a racemic mixture at a dose of 2.0 mg/kg (final concentration 2.0 mg/ml; Laboratorio Chile, Santiago, Chile). The vehicle solution consisted of 90% 0.9% NaCl (Sanderson S.A. Laboratory, Santiago, Chile), 5% ethanol (95% ethanol; Winkler Limitada, Santiago, Chile), and 5% Kolliphor (Sigma, Germany). The GRK2/3 inhibitor Cmpd101 was administered at a dose of 1.0 mg/kg (final concentration 1.0 mg/ml; Hello Bio, Bristol, United Kingdom). All treatments were delivered via intraperitoneal (i.p.) injection.

### 2.3. Experimental protocol

The experimental protocol consisted of handling, locomotor sensitization protocol, brain extraction, protein quantification, Western Blot, and statistical analysis. Locomotor activity was recorded using a Logitech HD C270 WebCam. Animals were randomly assigned to one of the following experimental groups: saline and vehicle (Sal-Veh; n = 4), saline and Cmpd101 (Sal-Cmpd; n = 4), acute AMPH and vehicle (AMPH(A)-Veh; n = 5), acute AMPH and Cmpd101 (AMPH(A)-Cmpd; n = 6), repeated AMPH and vehicle (AMPH(R)-Veh; n = 10), and repeated AMPH and Cmpd101 (AMPH(R)-Cmpd; n = 9).

### 2.4. AMPH locomotor sensitization

The AMPH-induced locomotor sensitization process was adapted following the protocol previously described by Gatica et al., 2020 and this is shown in Figure 1. This protocol begins during the light phase between 9:00 and 10:00 a.m. (Chilean time). Locomotor activity was measured in acrylic boxes (45 cm x 23 cm x 15.5 cm) and analyzed using ANY-maze software (Stoelting Co.). During the first two days, the animal’s baseline locomotor activity was evaluated in the locomotor activity box for one hour, after which an i.p. saline injection was administered and its activity in the box was recorded again for one hour.

**Figure 1.**
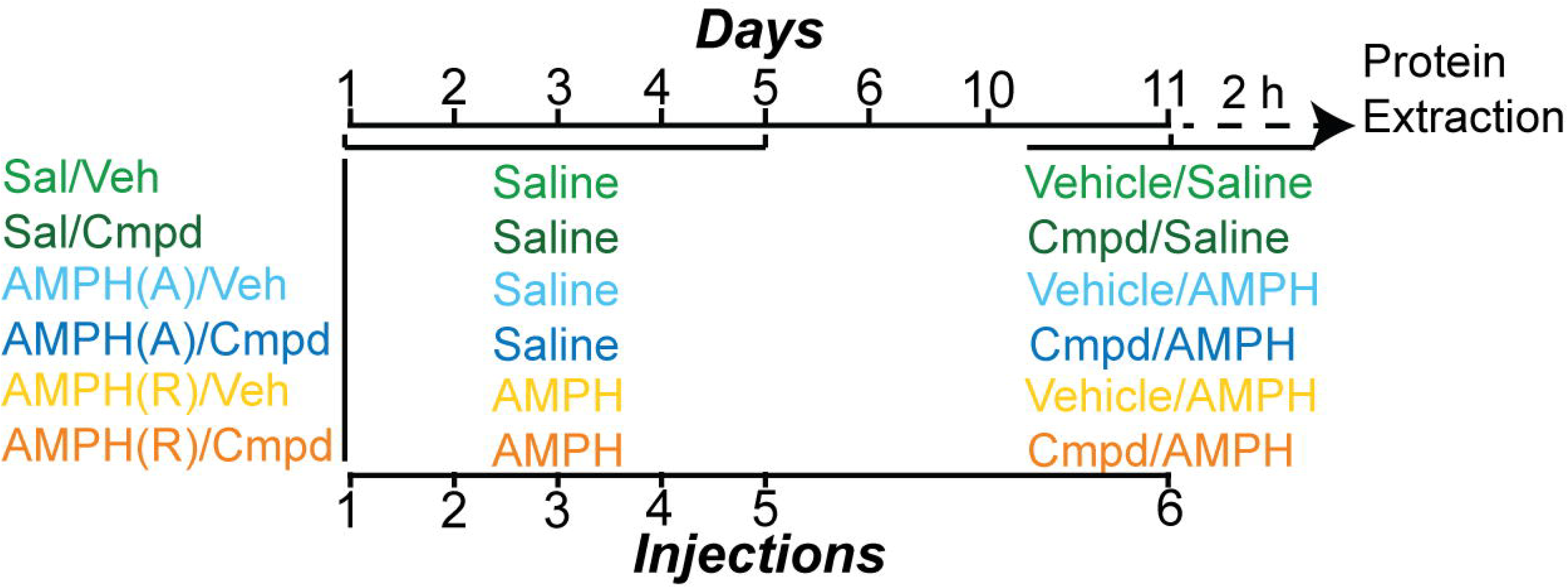
Experimental protocol

Locomotor sensitization was assessed by recording activity in the locomotor activity chamber for one hour at baseline, followed by i.p. injection of saline or AMPH, after which locomotion was recorded for an additional one and a half hours. This protocol was performed for five consecutive days. This was followed by a four-day period of withdrawal. Next, after measuring baseline locomotor activity for 1 hour, all groups were injected with saline solution i.p. and their locomotion was measured for an additional hour. On the last day, basal locomotor activity was again measured in the locomotor activity chamber for one hour, and then the rats (both the saline and AMPH groups) were injected with vehicle or Cmpd101 i.p. After the injection, locomotor activity was measured for 30 min. Then, all groups were injected i.p. with either saline or AMPH, and locomotor activity was measured for an hour and a half.

The AMPH Locomotion Score was calculated according to the formula by Gatica et al., 2020:

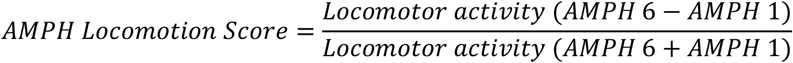

Rats with an AMPH locomotion score higher than 0.1 were considered sensitized, whereas rats with an AMPH locomotion score lower than 0.1 were considered non-sensitized.

On the other hand, the locomotor activity observed after injection 6 was normalized with respect to the previous injection (injection 5), calculating the percentage change using the formula:

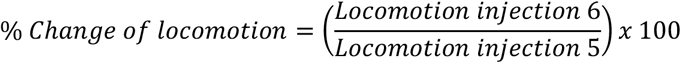

### 2.5. Protein extraction and quantification

Two hours after Cmpd101 or vehicle injection, rats were euthanized and their brains were extracted. Isoflurane 4% (Baxter, United States) was used as an anesthetic. When the animal achieved anesthetic plane (no reaction when squeezing the tail, hind leg, and interdigital space), a guillotine was used to obtain the head. Dissections of the brain regions were performed in a Petri dish on ice.

For total protein levels studies, the dorsomedial striatum (DMS) and dorsolateral striatum (DLS) were extracted. Brain tissue was homogenized directly in 2X Laemmli buffer (5% w/v SDS, 25% v/v glycerol, 150mM Tris-HCl pH 6.8) diluted to 1X with triple-distilled water. Subsequently, samples were boiled for 5 minutes at 80°C. Finally, ultrasonication was performed for 1 minute.

For synaptosome studies, the nucleus accumbens (NAc) and dorsal striatum (DS) were extracted. Tissue was extracted according to the method of Massaro Tieze et al., 2022. The brain tissue was homogenized in 10 mM HEPES solution with 320 mM sucrose at pH 7.4. The samples were homogenized using a glass–Teflon homogenizer. Subsequently, they were centrifuged (800 G at 4°C for 10 minutes). The supernatant was transferred to another Eppendorf tube and centrifuged again at 9000 G at 4°C for 15 minutes. The supernatant was discarded and 500 µL of HEPES buffer was added to the pellet. It was centrifuged again at 9000 G at 4°C for 15 minutes. The supernatant was discarded, and the pellet was resuspended with 100 μL of HEPES buffer.

The proteins obtained from each extraction were quantified using the micro-BCA Protein Assay Kit (Pierce-Thermo Scientific).

### 2.6. Western blot

Proteins extracts were separated on an 8% SDS–PAGE gel and subsequently transferred to a 0.2 µm nitrocellulose membrane (Bio-Rad Laboratories) for 2 h at 400 mA and 4 °C. Following transfer, membranes were washed once with TBS for 5 min and incubated in Ponceau red solution (0.2% Ponceau red in 5% acetic acid) for 15 min. Membranes were then washed with distilled water, and images of Ponceau red staining were acquired using a photodocumentation system (GeneGnome XRQ, Syngene). The membrane was then cut according to molecular weight to separate GRK2 and GRK5 from D2R. Subsequently, membranes were incubated in blocking solution (5% non-fat dry milk diluted in TBS–Tween 0.1%) for 1 h. Membranes were then incubated overnight at 4 °C with the following primary antibodies diluted in TBS–Tween 0.1%: mouse anti-GRK2 (1:5000; sc-13143, Santa Cruz Biotechnology) and mouse anti-GRK5 (1:5000; sc-518005, Santa Cruz Biotechnology) for membranes containing proteins >55 kDa, and mouse anti-D2R (1:1000; sc-13143, Santa Cruz Biotechnology) for membranes containing proteins <55 kDa. To optimize signal detection, antibody concentrations were adjusted as follows: in the DS (syn) region, the mouse anti-GRK5 antibody was used at a dilution of 1:1000, whereas in the NAc (syn) region, mouse anti-GRK2 and mouse anti-GRK5 antibodies were used at dilutions of 1:2500 and 1:1000, respectively. After incubation, membranes were washed with 0.1% Tween in TBS and incubated with TBS-Tween 0.1% anti-mouse horseradish peroxidase (HRP) donkey (1:5000; 715-035-150, Jackson ImmunoResearch Laboratories) diluted in TBS-Tween 0.1% and incubated for 1 h at room temperature. Membranes were subsequently incubated with SuperSignal West Pico chemiluminescent substrate (ThermoFisher Scientific, USA) and imaged with a GeneGnome photodocumentation system. Protein bands corresponding to GRK2, GRK5, and D2R were quantified using ImageJ software (National Institutes of Health, USA). Band optical density values were obtained with ImageJ and normalized to bands of equal molecular weight detected by Ponceau red staining, a validated total protein loading control (Bettencourta et al., 2020, Wang et al., 2022).

### 2.7. Statistical analysis

All data processing and statistical analyses were performed using Python (version 3.12.3), primarily utilizing the statsmodels and pandas libraries. To avoid the mathematical artifacts and non-linearities often introduced by simple ratio-based normalization, total protein loading control (Ponceau S) was incorporated into all models as a continuous covariate rather than a direct denominator. To analyze main effects and categorical group differences across brain regions while rigorously controlling for technical batch variability, we employed Linear Mixed-Effects Models (LMM), similar to Omondi et al., 2024. The experimental treatment was modeled as a fixed effect, Ponceau S optical density as a fixed covariate, and the Western blot membrane identifier was assigned as a random intercept. This approach isolates true biological variance by explicitly modeling the baseline luminescence differences between individual membranes. Group visualizations and pairwise comparisons were derived from the Estimated Marginal Means (EMMs) generated by the LMM, with variance reported as the model-adjusted 95% Confidence Intervals (95% CI). To evaluate the continuous interaction between the subjects’ phenotypic behavior (AMPH locomotion score) and the pharmacological treatment, we utilized a sequential robust regression pipeline.

Because crossing a continuous gradient with a categorical variable alongside a random effect often leads to singular fit or convergence failures in standard LMM algorithms, technical noise was handled sequentially. First, the variance attributable to the analytical batch (membrane effect) was statistically isolated and subtracted from the raw target protein densities, yielding a residualized, batch-cleaned protein variable. Second, an Ordinary Least Squares (OLS) regression was fitted to this cleaned data, modeling the interaction between the AMPH score and Treatment. Crucially, the model employed Heteroskedasticity-Consistent type 3 (HC3) robust standard errors. The use of HC3 estimators protects the model against unequal variances across the continuous spectrum, preventing inflated Type I error rates.

Mixed model ANOVA was used when required. The alpha level for statistical significance was set a priori at p < 0.05. To control for the False Discovery Rate (FDR) during multiple testing scenarios, p-values were adjusted using the Benjamini-Hochberg procedure.

Full statistical reports are available in the Supplementary Material.

## 3 Results

### 3.1 Effect of Cmpd101 on basal and injection-induced locomotor activity

To assess the effect of Cmpd101 on locomotor activity under basal conditions and following repeated AMPH injection, we compared activity levels across experimental groups (Fig. 2). The experimental protocol is illustrated in Fig. 2A, where basal locomotor activity was recorded for 30 min prior to injection (Veh or Cmpd101), followed by a 30 min post-injection recording period. The Saline and Acute AMPH groups were combined, as both received only saline injections until this point in the protocol. A linear mixed model revealed a significant interaction between stage (basal vs injection) and group (Sal vs AMPH(R)) (p = 0.015; see Supplementary Material for more details), indicating that the effect of injection on locomotor activity differed between saline- and AMPH(R)-treated animals. Specifically, saline-treated animals showed a reduction in locomotor activity following injection, whereas AMPH(R)-treated animals exhibited an opposite trend. No significant interaction involving the test drug (Cmpd101 vs Veh) was observed.

**Figure 2.**
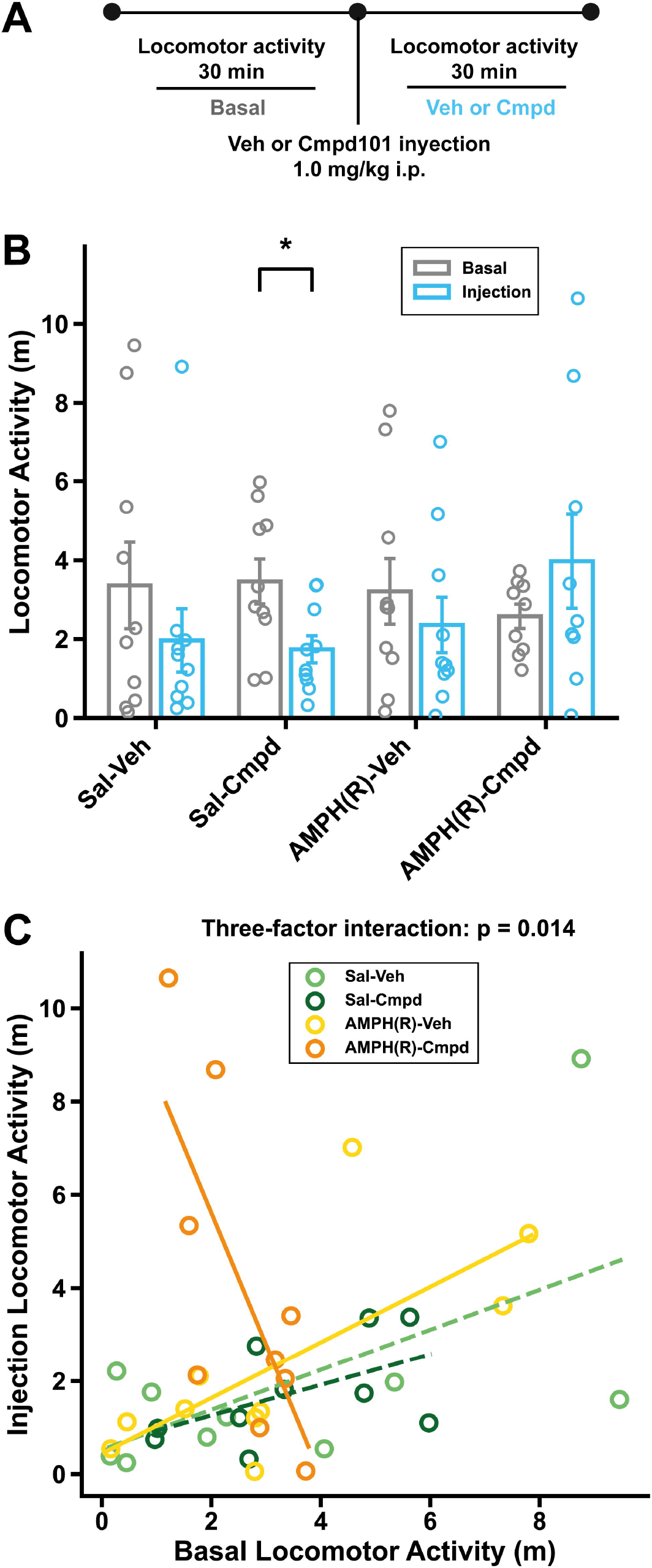
Effect of Cmpd101 on basal and injection-induced locomotor activity. A) Locomotor activity measured under basal conditions and following injection across treatment groups: saline and vehicle (Sal-Veh; n = 10), saline and Cmpd101 (Sal-Cmpd; n = 9), repeated AMPH and vehicle (AMPH(R)-Veh; n = 10), and repeated AMPH and Cmpd101 (AMPH(R)-Cmpd; n = 9). Bars represent mean ± SEM and circles represent individual data points. A significant difference between conditions is indicated (**p < 0.01). B) Relationship between basal and injection-induced locomotor activity across treatment groups. Each point represents an individual animal. Linear regression analysis shows a significant three-factor interaction (p = 0.014), indicating that the relationship between basal and injection locomotor activity differs across pre-treatment conditions.

To further examine the relationship between basal and injection-induced locomotor activity, we performed a linear regression analysis across groups (Fig. 2C). This analysis revealed a significant three-factor interaction between treatment, inhibitor, and basal activity (p = 0.014), indicating that the relationship between basal and injection-induced locomotion differs depending on experimental condition. Notably, Cmpd101 treatment altered the slope of this relationship, particularly in AMPH(R) animals: rats with high basal locomotion in the AMPH(R) group showed lower locomotor activity after Cmpd101 injection than those with low basal locomotion. These results suggest that the inhibitor modifies the association between basal locomotor activity and injection-induced locomotor responses in rats repeatedly treated with AMPH.

### 3.2 Cmpd101 does not alter AMPH-induced locomotor activity nor the expression of locomotor sensitization

To determine whether Cmpd101 modifies AMPH-induced locomotor activity, we first compared the locomotor response to the sixth injection across experimental groups (Fig. 3A; timeline in Fig. 1). As expected, animals exposed to AMPH (both acute and repeated) displayed higher locomotor activity than saline-treated controls (Fig. 3, see Supplementary Material for more details). In contrast, no differences were observed between vehicle- and Cmpd101-treated animals within the same treatment condition, indicating that the inhibitor did not alter the locomotor response to acute or repeated AMPH injection (Fig. 3A).

**Figure 3.**
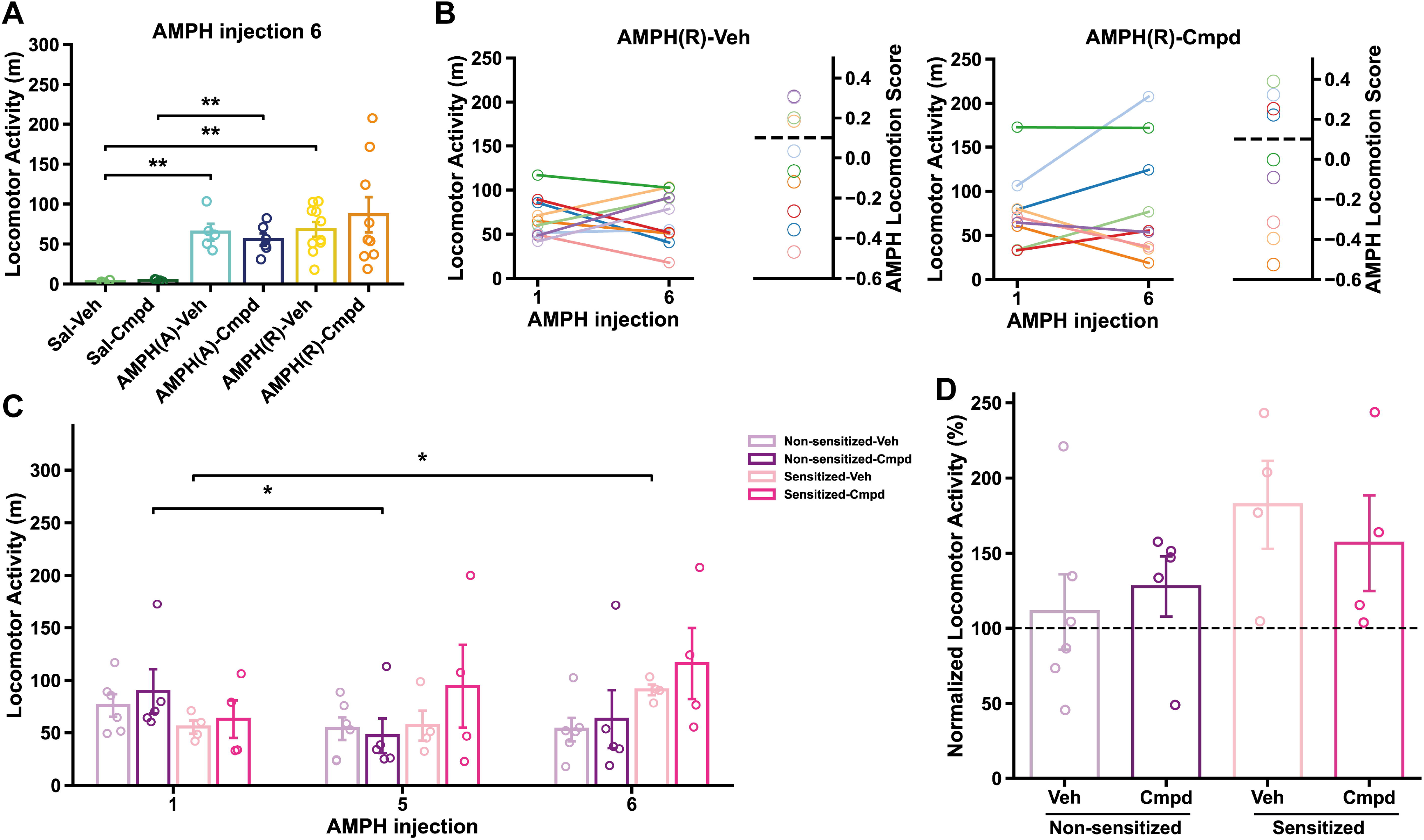
Cmpd101 did not modify the expression of AMPH locomotor sensitization. A) Locomotor activity on injection 6 across treatment groups: saline and vehicle (Sal-Veh; n = 4), saline and Cmpd101 (Sal-Cmpd; n = 4), acute AMPH and vehicle (AMPH(A)-Veh; n = 5), acute AMPH and Cmpd101 (AMPH(A)-Cmpd; n = 6), repeated AMPH and vehicle (AMPH(R)-Veh; n = 10), and repeated AMPH and Cmpd101 (AMPH(R)-Cmpd; n = 9). ** p< 0.01 using Benjamini-Hochberg post test. Bars represent mean ± SEM and circles represent individual data points. B) Individual locomotor activity trajectories on injection 1 and 6 in AMPH(R)-Veh and AMPH(R)-Cmpd groups (left panels), and corresponding AMPH Locomotion Scores (right panels). Each line and matching circle represents the same animal. Dashed line indicates the threshold used to classify rats as sensitized (AMPH Locomotion Score ≥ 0.1) or non-sensitized (< 0.1). C) Time course of locomotor activity (days 1, 5, and 7) in rats classified as sensitized (n = 4 for AMPH(R)-Veh and AMPH(R)- Cmpd) and non-sensitized (n = 6 for AMPH(R)-Veh and n = 5 for AMPH(R)-Cmpd), treated with vehicle or Cmpd101. * p< 0.05 using Benjamini-Hochberg post test. Bars represent mean ± SEM and circles represent individual data points. D) Normalized locomotor activity (injection 6/injection 5) in sensitized and non-sensitized rats under vehicle and Cmpd101 conditions. Bars represent mean ± SEM and circles represent individual data points.

We then analyzed repeated AMPH-treated animals separately in the AMPH(R)-Veh and AMPH(R)- Cmpd groups by comparing locomotor activity between injections 1 and 6 and calculating the AMPH Locomotion Score (Gatica et al., 2020, 2022; Fig. 3B). In both groups, individual responses were heterogeneous, with some animals showing increased locomotion at injection 6 relative to injection 1 (sensitized) and others showing little change or a reduction (non-sensitized). The proportion of animals that sensitized and did not sensitize to AMPH in the AMPH(R)-Veh and AMPH(R)-Cmpd groups was similar. Specifically, 40% of animals were classified as sensitized in the AMPH(R)-Veh group and 44.4% in the AMPH(R)-Cmpd group, whereas 60% and 55.6% were non-sensitized, respectively. Consistent with this, comparison of AMPH Locomotion Score mean values between groups did not reveal significant differences (unpaired t-test, p=0.9524).

To further examine this variability, AMPH(R) animals were classified as non-sensitized or sensitized, and their locomotor activity was compared across injections 1, 5, and 6 (Fig. 3C). As expected, sensitized animals exhibited higher locomotor activity than non-sensitized animals in the sixth AMPH injection, consistent with the development of locomotor sensitization. However, no differences were detected between vehicle- and Cmpd101-treated animals in either the non-sensitized or the sensitized groups (for more statistical details, see Supplementary Material).

Finally, normalization of locomotor activity (injection 6/injection 5) confirmed a trend of higher locomotor activity in sensitized compared with non-sensitized animals both in vehicle and Cmpd101 groups. In the sensitized animals, Cmpd101 treatment did not significantly modify this effect (Fig. 3D).

Together, these results indicate that Cmpd101 did not significantly alter acute AMPH-induced hyperlocomotion or the expression of locomotor sensitization.

### 3.3 Cmpd101 treatment selectively modifies GRK levels in the DMS and NAc of AMPH-treated rats

We next assessed the effect of Cmpd101 on protein levels of D2R, GRK2, and GRK5 across four striatal regions (DLS, DMS, DS synaptosomal fraction, and NAc synaptosomal fraction) (Fig. 4, see Supplementary Figure 1 for representative western blots images). To distinguish between global protein levels and localized synaptic adaptations, biochemical analyses were performed using both total tissue homogenates and enriched synaptosomal fractions. While total tissue levels provide a broad measure of protein abundance, the isolation of synaptosomes allows for a more precise evaluation of the protein pools actively involved in neurotransmission and receptor trafficking at the synapse. Protein levels were analyzed using linear mixed models (LMM), which allow the inclusion of loading controls as covariates and the modeling of technical sources of variability, such as differences across gels and replicates, as random effects. This approach improves the accuracy and reproducibility of Western blot quantification by accounting for both biological and technical variability (Omondi et al., 2024).

**Figure 4.**
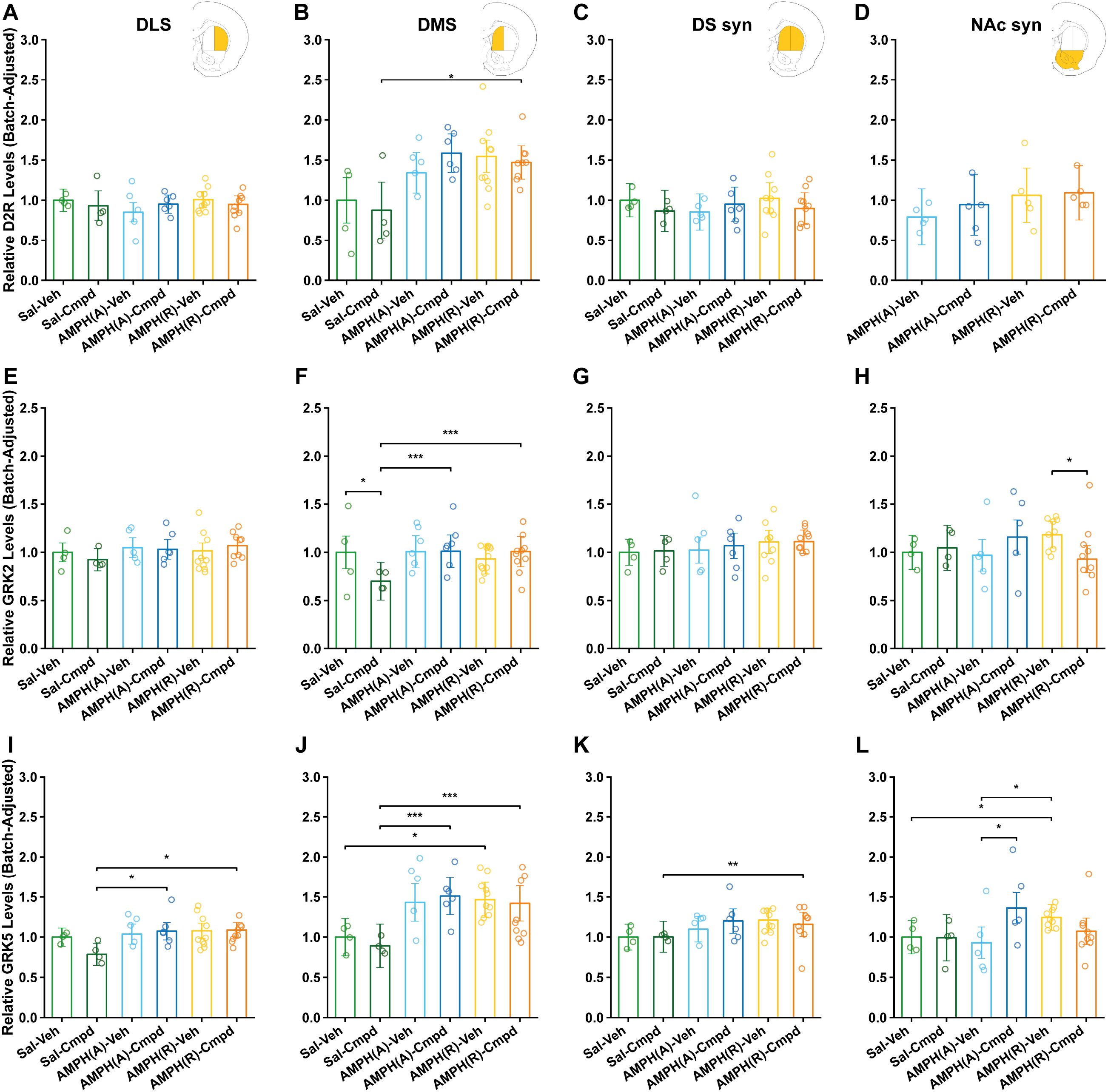
Cmpd101 induces region-specific changes in GRK levels in AMPH-treated rats. A-D) Relative D2R protein levels (batch-adjusted) on: A) dorsal lateral striatum (DLS), B) dorsal medial striatum (DMS), C) dorsal striatum synaptosomal fraction (DS syn), and D) nucleus accumbens synaptosomal fraction (NAc syn). Treatment groups: saline and vehicle (Sal-Veh; n = 4), saline and Cmpd101 (Sal-Cmpd; n = 4), acute AMPH and vehicle (AMPH(A)-Veh; n = 5), acute AMPH and Cmpd101 (AMPH(A)-Cmpd; n = 6), repeated AMPH and vehicle (AMPH(R)-Veh; n = 10), and repeated AMPH and Cmpd101 (AMPH(R)-Cmpd; n = 9). E-H) Relative GRK2 protein levels (batch-adjusted) across the same regions and treatment groups. I-L) Relative GRK5 protein levels (batch-adjusted) across the same regions and treatment groups. Bars represent Estimated Marginal Means (EMM) and confidence interval 95%, and circles represent individual data points. Statistical comparisons were performed using linear mixed-effects models, and significant effects are indicated as *p < 0.05, **p < 0.01, ***p < 0.001 using Benjamini-Hochberg post test.

D2R levels was not significantly altered by Cmpd101 treatment in any region or experimental group, as no consistent differences were observed between vehicle- and Cmpd101-treated animals (Fig. 4A-D). For GRK2, Cmpd101 treatment induced selective, region- and group-specific effects (Fig. 4E-H). In the DMS, GRK2 levels were significantly reduced in the Sal-Cmpd group compared to its corresponding vehicle control and the AMPH(A)-Cmpd and AMPH(R)-Cmpd groups (Fig. 4F). A decrease was observed in the NAc synaptosomal fraction in AMPH(R)-Cmpd animals relative to AMPH(R)-Veh (Fig. 4H). No consistent differences were detected in the remaining regions or treatment conditions. GRK5, a member of the GRK4 subfamily, was measured to control for changes in a GRK subfamily distinct from GRK2/3. GRK5 levels showed region-dependent variations (Fig. 4I-L). Notably, a significant increase in GRK5 levels was observed in the NAc synaptosomal fraction in the AMPH(A)-Cmpd group compared to AMPH(A)-Veh, indicating an effect of Cmpd101 under acute AMPH conditions. However, no consistent pattern of change was observed between vehicle- and Cmpd101-treated animals across all regions. Instead, most of the differences in GRK5 levels were observed between experimental groups (Sal vs AMPH, both acute and repeated), suggesting that AMPH exposure, rather than Cmpd101 treatment independently, is the main driver of GRK5 alterations. For instance, in some regions, GRK5 levels were increased in AMPH-treated animals compared to saline controls within the same treatment condition (for more statistical details, see Supplementary Material).

Overall, these results indicate that while Cmpd101 does not affect D2R levels, it selectively reduces GRK2 levels in specific regions and experimental groups. In contrast, GRK5 levels showed predominantly AMPH-dependent changes, with limited effects of Cmpd101, except for a significant increase observed in the NAc synaptosomal fraction under acute AMPH conditions.

### 3.4 Cmpd101 modulates protein-behavior association in a region-specific manner

To examine whether protein levels depend on the association between Cmpd101 effect and the degree of locomotor sensitization, we used an OLS+HC3 regression analysis with AMPH Locomotion Score as the predictor variable to test for an interaction with the treatment (Fig. 5).

**Figure 5.**
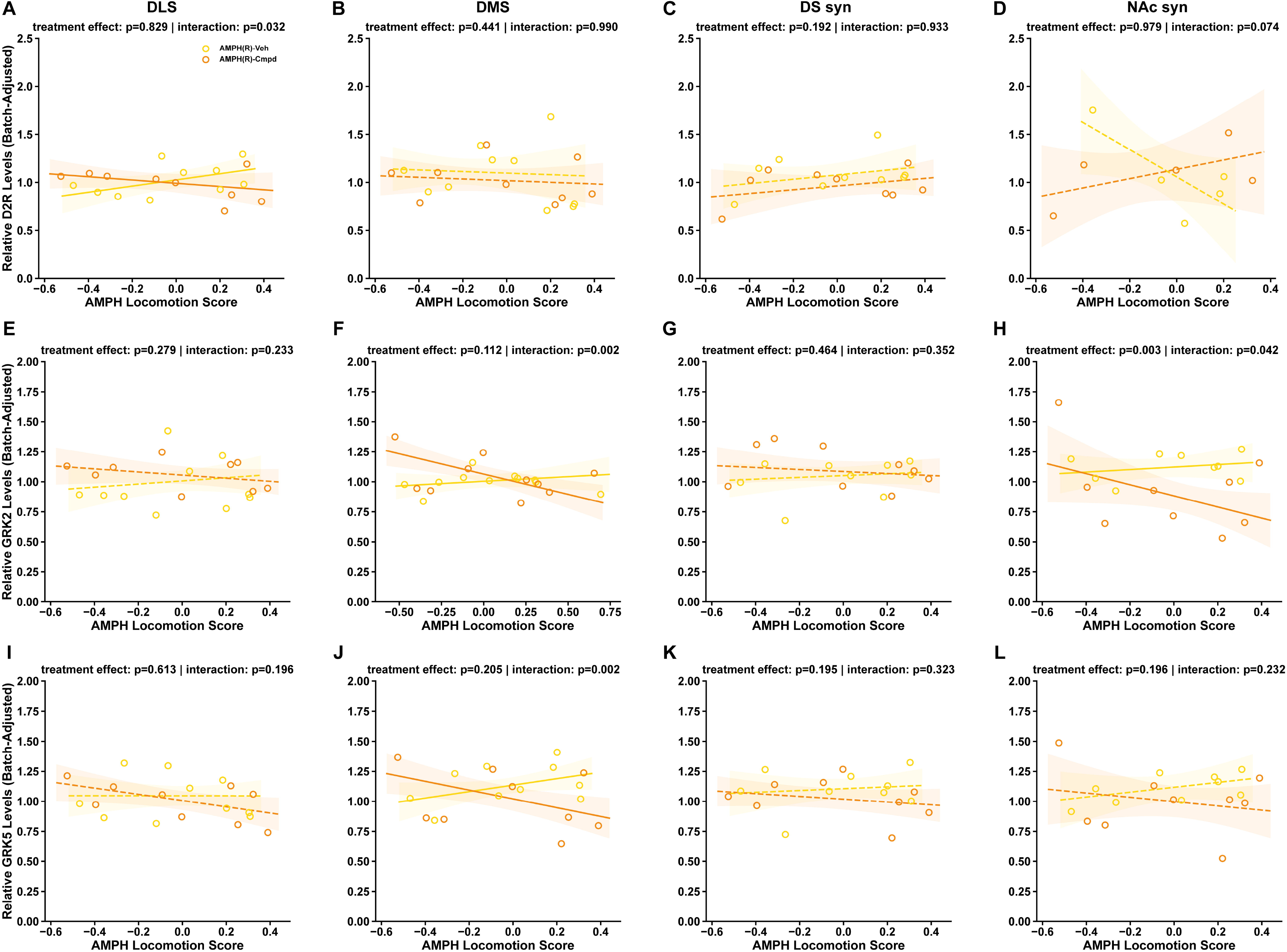
Cmpd101 changes D2R, GRK2, and GRK5 protein levels is locomotor sensitization dependent. A-D) Relationship between AMPH Locomotion Score and relative D2R protein levels (batch-adjusted) in A) dorsal lateral striatum (DLS), B) dorsal medial striatum (DMS), C) dorsal striatum synaptosomal fraction (DS syn), and D) nucleus accumbens synaptosomal fraction (NAc syn) in repeated AMPH groups (AMPH(R)-Veh, n = 10; AMPH(R)-Cmpd, n = 9). E-H) Relationship between AMPH Locomotion Score and relative GRK2 protein levels (batch-adjusted) across the same regions and treatment groups. I-J) Relationship between AMPH Locomotion Score and relative GRK5 protein levels (batch-adjusted) across the same regions and treatment groups. Each point represents an individual animal. Lines represent fitted regression models and shaded areas indicate 95% confidence intervals. Statistical analyses were performed using linear models including treatment and interaction terms; reported p-values correspond to treatment effects and treatment × AMPH Locomotion Score interactions for each region.

For D2R, no significant treatment effects or interactions were observed in the DMS or DS synaptosomal fraction (Fig. 5B-C). In contrast, a significant interaction was detected in the DLS (interaction p = 0.032; Fig. 5A, see Supplementary Material for more details), indicating that the relationship between D2R levels and AMPH Locomotion Score differed between vehicle- and Cmpd-treated animals. A similar trend was observed in the NAc synaptosomal fraction, although it did not reach statistical significance (Fig. 5D). For GRK2, a significant interaction between treatment and locomotion score was observed in the DMS (interaction p = 0.002; Fig. 5F), indicating that Cmpd101 altered the relationship between GRK2 levels and the AMPH Locomotion Score. Additionally, in the NAc synaptosomal fraction, both a significant treatment effect (p = 0.003) and interaction (p = 0.042) were detected (Fig. 5H), supporting a region-specific modulation of GRK2-behavior relationships by Cmpd101. No significant effects were observed in the DLS or DS synaptosomal fraction (Fig. 5E, G). For GRK5, a significant interaction was observed in the DMS (interaction p = 0.002; Fig. 5J), while no significant effects were detected in the other regions (Fig. 5I, K-L). (for more statistical details, see Supplementary Material).

Overall, these results indicate that Cmpd101 does not uniformly alter protein levels but instead modulates the relationship between specific proteins and locomotor sensitization in a region-dependent manner, particularly for D2R in the DLS, GRK2 and GRK5 in the DMS and GRK2 in the NAc.

## 4. Discussion

Susceptibility to psychostimulant-induced sensitization is associated with individual differences in dopaminergic function (Gatica et al., 2020; Marinelli & White, 2000; Nader et al., 2006; Piazza et al., 1989). The present study provides evidence of individual variability in AMPH-induced locomotor sensitization and its association with region-specific modulation of protein-behavior relationships. Consistent with previous literature, repeated AMPH exposure induced locomotor sensitization in a subset of animals, reflecting inter-individual vulnerability (Casanova et al., 2013; Pierce & Kalivas, 1997). Within the framework of the incentive-sensitization theory, this progressive hypersensitivity reflects a pathological usurpation of reward-related learning, or ’wanting’ (Robinson & Berridge, 2025). In our data, this variability was also evident in the heterogeneous AMPH Locomotion Scores.

The fact that not all animals developed sensitization aligns with the existing literature, which highlights marked individual differences in vulnerability to psychostimulant addiction, often predicted by baseline behavioral endophenotypes such as responses to novelty or impulsivity (Belin & Deroche-Gamonet, 2012; Egervari et al., 2018; Piazza & Deroche-Gamonet, 2013).

Our behavioral results show that Cmpd101 produced opposite trends in locomotor activity in saline-versus AMPH(R)-treated animals, supporting a state-dependent effect of GRK2/3 inhibition, likely reflecting the dual role of D2 receptors. Presynaptically, D2 autoreceptors inhibit dopamine release, whereas postsynaptically, they modulate neuronal excitability. Thus, GRK2/3 inhibition may enhance autoreceptor-mediated negative feedback in basal conditions but favor postsynaptic signaling in a sensitized dopaminergic state, as previously proposed (Calipari et al., 2014; Mandeville et al., 2023). Despite this, Cmpd101 did not significantly modify AMPH-induced locomotor activity or the proportion of sensitized animals, suggesting that acute GRK2/3 inhibition is insufficient to alter the expression of locomotor sensitization. Genetic models demonstrate that while selectively deleting GRK2 in D2R-expressing neurons reduces psychostimulant sensitization (Daigle et al., 2014), global GRK3 ablation exacerbates AMPH-induced hyperlocomotion (Sellgren et al., 2021). Therefore, the simultaneous pharmacological inhibition of both GRK2 and GRK3 by Cmpd101 likely engages competing neuroadaptations that cancel each other out, yielding no net change in behavioral output.

We show that GRK2/3 inhibition did not alter total D2R levels in any group. This contrasts with previous in vitro work showing that Cmpd101 can interfere with D2R internalization in cultured cells (Sánchez-Soto et al., 2023), indicating that blocking receptor desensitization at the cellular level does not necessarily translate into gross protein changes or behavioral effects under these in vivo conditions. Furthermore, while imaging studies often report decreased D2R availability following chronic drug exposure (Everitt et al., 2008; Nader et al., 2006), those methodologies measure surface binding rather than total protein content. Thus, drug-induced adaptations likely involve subcellular redistribution or functional uncoupling, altering surface availability without changing gross protein levels, suggesting that GRK-mediated regulation here is functional rather than structural.

In contrast, we observed that Cmpd101 induced selective and region-dependent changes in GRK2 levels. A reduction in DMS was observed under saline conditions and in the NAc synaptosomal fraction of AMPH(R) rats, as evidenced by decreased GRK2 levels and an interaction between GRK2 levels and AMPH Locomotion Score. The NAc is critically involved in reward and incentive salience (Everitt & Robbins, 2016; Robinson & Berridge, 1993) and is the primary focus of the early binge/intoxication stage of addiction, where it integrates glutamatergic inputs from the prefrontal cortex and amygdala, and receives dense dopaminergic innervation from the ventral tegmental area (Egervari et al., 2018; Koob & Volkow, 2010). The specific interaction between AMPH Locomotion Score and Cmpd101 on NAc synaptic GRK2 highlights this region’s extreme sensitivity to dopaminergic overflow and subsequent receptor regulatory mechanisms (Anderson & Hearing, 2019; Jentsch & Pennington, 2014). Moreover, the detection of this effect in the synaptosomal fraction suggests localized synaptic regulation, highlighting that systemic administration of Cmpd101 can produce region and compartment-specific effects. Alterations in NAc GRK2 signaling may disrupt the balance between D1 (direct) and D2 (indirect) medium spiny neuron pathways, a fundamental neuroadaptation required for the expression of locomotor sensitization (Egervari et al., 2018; Philibin et al., 2011).

Regarding GRK5, its levels appeared to be primarily driven by AMPH exposure rather than by inhibition of GRK2/GRK3. This aligns with broader literature indicating that psychostimulants can differentially regulate distinct GRK isoforms; for instance, chronic cocaine has been shown to selectively upregulate GRK5 mRNA in specific limbic regions, while downregulating GRK2, GRK3, and GRK5, but not GRK6 protein levels in the prefrontal cortex of cocaine addicts (Álvaro-Bartolomé & García-Sevilla, 2013)

An important observation is the lack of a clear relationship between GRK alterations and D2R levels. Although classical *in vitro* studies have shown that GRK2 regulates D2R phosphorylation and internalization (Iwata et al., 1999; Kim et al., 2001), the changes observed here in GRK2 or GRK5 were not accompanied by parallel changes in total D2R levels. This dissociation aligns with findings demonstrating that GRK-mediated phosphorylation is not strictly required for D2R internalization but rather regulates its post-endocytic recycling (Cho et al., 2010; Namkung et al., 2009b), and that GRK2 can constitutively modulate D2R levels through phosphorylation-independent mechanisms (Namkung et al., 2009a). Furthermore, the lack of parallel changes highlights the complex, homeostatic maintenance of D2R levles in vivo, which may not linearly reflect the GRK-driven dynamics observed in heterologous overexpression models.

A key finding is the regional specificity of protein alterations. Protein levels were mainly changed in the DMS and NAc rather than the DLS in rats with repeated AMPH administration. This dissociation is relevant to the current understanding of the neurocircuitry of addiction. The transition from initial, recreational drug use to compulsive taking is mediated by a functional shift from the NAc to the dorsal striatum. Within the dorsal striatum, the DMS is primarily involved in action-outcome (goal-directed) learning, whereas the DLS mediates stimulus-response (habitual) behaviors (Cataldi et al., 2022; Everitt & Robbins, 2013, 2016; Yin & Knowlton, 2006). The alterations in the DMS, rather than the DLS, suggest that the repeated AMPH protocol captures an early to intermediate stage of neuroplasticity where behavior is still largely goal-directed, prior to the full consolidation of compulsive habits. The selective effects of Cmpd101 on GRK2 in the NAc and DMS further support the idea that GRK-dependent signaling contributes differentially to distinct stages of addiction, with NAc alterations being particularly sensitive during early motivational processes and reward-related learning.

We found a dissociation between the molecular alterations and behavioral outcomes after repeated AMPH administration. While Cmpd101 altered specific protein levels in a region-dependent manner, it did not significantly reduce the expression of locomotor sensitization. This may reflect compensatory mechanisms within the dopaminergic system or the recruitment of D1R and glutamatergic pathways (Egervari et al., 2018; Philibin et al., 2011). The observed interactions between AMPH Locomotion Score and D2R levels in the DLS, GRK2 levels in the DMS/NAc, and GRK5 levels in the DMS, suggest that GRK2/3 inhibition modifies the relationship between the degree of AMPH locomotor sensitization and protein levels in these regions. This is particularly relevant in the context of addiction, where individual variability plays a major role (Anthony et al., 1994; Egervari et al., 2018; Lopez-Quintero et al., 2011). Examining these effects in paradigms such as conditioned place preference (CPP) or self-administration will be critical to determine whether this modulation affects rewarding properties more than gross locomotor output.

While the present study provides critical insights into the regional neuroadaptations associated with AMPH sensitization, it is important to note that the pharmacological manipulation of GRK2/3 was restricted to a single acute systemic administration of Cmpd101 at a dose of 1.0 mg/kg. While this acute intervention provides an initial proof of concept, it may not be enough to counteract the deep and long-lasting neuroplastic changes induced by repeated AMPH exposure. GRKs exhibit highly dynamic, temporally regulated alterations in response to chronic psychostimulant administration.

Therefore, a single acute dose limits the ability to detect broader behavioral and biochemical effects that might require prolonged inhibition of the target. Future studies should therefore include dose-response analyses and chronic Cmpd101 administration protocols aligned with the temporal profile of repeated AMPH exposure.

In summary, GRK2/3 inhibition by Cmpd101 produces region-specific molecular effects and reshapes protein-behavior relationships without significantly altering locomotor sensitization. These findings support a model in which GRKs act as modulators of dopaminergic signaling in a context- and region-dependent manner, rather than as direct determinants of behavioral output.

## Data availability statement

The raw data supporting the conclusions of this article will be made available by the authors, without undue reservation.

## Ethics statement

The animal study was approved by Comité Etico Científico en Cuidado Animal y Ambiente de la Pontificia Universidad Católica de Chile. The study was conducted in accordance with the local legislation and institutional requirements.

## Author contributions

CS: Writing – Original draft, Writing – Review & editing, Investigation, Data curation, Methodology, Formal Analysis, Visualization; MEA: Writing – Review & editing, Conceptualization, Supervision, Funding acquisition, Resources; RIG: Writing – Original draft, Writing – Review & editing, Conceptualization, Investigation, Software, Data curation, Methodology, Supervision, Formal Analysis, Project administration, Validation, Funding acquisition, Resources, Visualization.

## Generative AI Statement

During the preparation of this work, the authors used Gemini and Grammarly in order to improve the clarity and grammar of the English text. After using this tool, the authors reviewed and edited the content as needed and take full responsibility for the content of the publication.

