## Supplementary Material for "Dissociation of Molecular and Behavioral Neuroadaptations Following Acute GRK2/3 Inhibition in Amphetamine-Treated Rats"

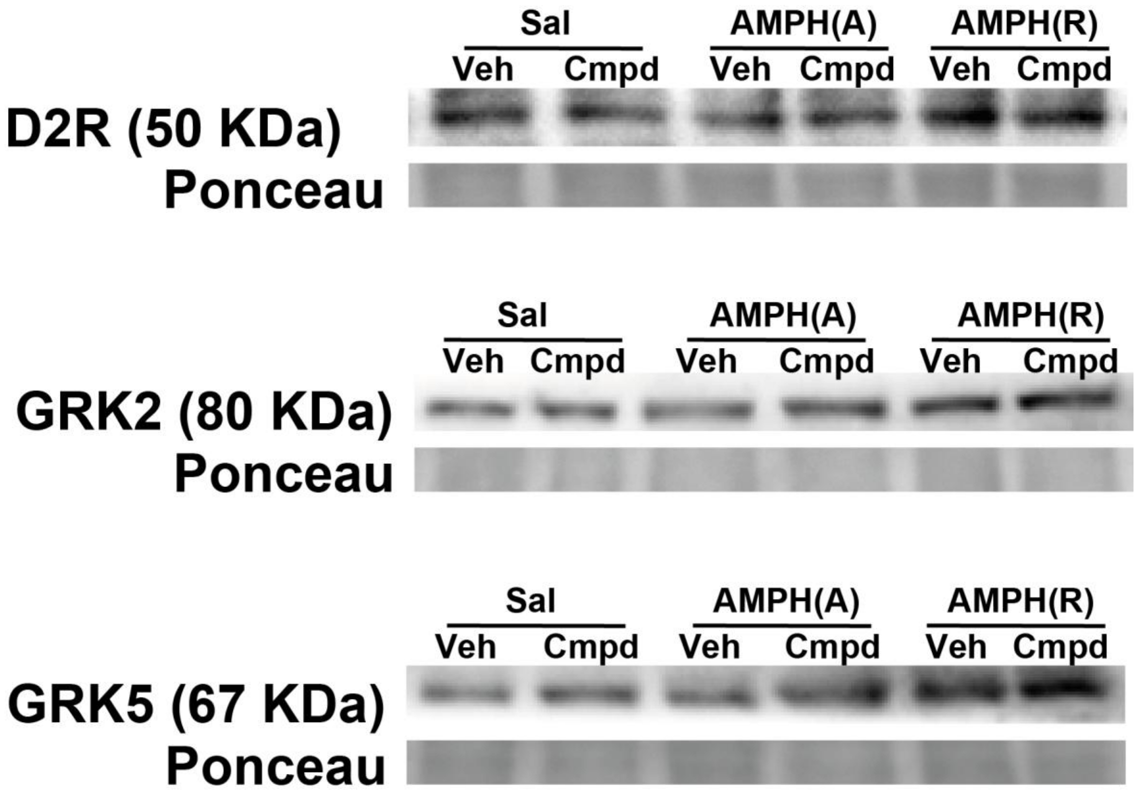

**Supplementary figure 1.** Representative Western blot images of D2R, GRK2, and GRK5 across the six experimental groups.

**Table 1**. Statistical analysis of Fig. 2B.

| 2x2x2 Linear Mixed Model (Repeated Measures) | |
|---|---|
| Model | Mixed LM |
| No. Observations | 78 |
| No. Groups | 39 |
| Min. group size | 2 |
| Max. group size | 2 |
| Mean group size | 2.0 |
| Dependent variable | Locomotor |
| Method | REML |
| Scale | 3.9201 |
| Log-Likelihood | -169.0195 |
| Coverged | Yes |

| | Coef | Std. Err. | z | p | 95% CI (Lower) | 95% CI (Upper) |
|---|---|---|---|---|---|---|
| Intercept | 2.578 | 0.818 | 3.150 | 0.002 | 0.974 | 4.182 |
| C(Stage_Label)[T.Injection] | 1.397 | 0.933 | 1.496 | 0.135 | -0.433 | 3.226 |

| | | | | | | |
|---|---|---|---|---|---|---|
| C(Pre_Treatment)[T.Sal] | 0.883 | 1.128 | 0.783 | 0.434 | -1.328 | 3.094 |
| C(Test_Drug)[T.Veh] | 0.632 | 1.128 | 0.561 | 0.575 | -1.579 | 2.843 |
| C(Stage_Label)[T.Injection]: C(Pre_Treatment)[T.Sal] | -3.118 | 1.287 | -2.423 | 0.015 | -5.639 | -0.596 |
| C(Stage_Label)[T.Injection]: C(Test_Drug)[T.Veh] | -2.248 | 1.287 | -1.748 | 0.081 | -4.770 | 0.273 |
| C(Pre_Treatment)[T.Sal]: C(Test_Drug)[T.Veh] | -0.734 | 1.574 | -0.467 | 0.641 | -3.820 | 2.351 |
| C(Stage_Label)[T.Injection]: C(Pre_Treatment)[T.Sal]: C(Test_Drug)[T.Veh] | 2.576 | 1.795 | 1.435 | 0.151 | -0.942 | 6.095 |
| Group Var | 2.107 | 0.694 | | | | |

**Table 2**. Statistical analysis of Fig. 2C.

| OLS Regression with HC3 (Injection ~ Basal_centered * Pre_Treatment * Test_Drug) | |
|---|---|
| Dep. Variable | Injection |
| R-squared | 0.528 |
| Model | OLS |
| Adj. R-squared | 0.422 |
| Method | Least Squares |
| F-statistic | 3.492 |
| Date | Sat, 11 Apr 2026 |
| Prob (F-statistic) | 0.00709 |
| Time | 17:39:02 |
| Log-Likelihood | -76.438 |
| No. Observations | 39 |
| AIC | 168.9 |
| Df Residuals | 31 |
| BIC | 182.2 |
| Df Model | 7 |
| Covariance Type | HC3 |

| Variable | Coef | Std. Error | z | p | 95% CI (Lower) | 95% CI (Upper) |
|---|---|---|---|---|---|---|
| Intercept | 2.3079 | 0.693 | 3.332 | 0.001 | 0.950 | 3.666 |
| C(Pre_Treatment)[T.Sal] | -0.6645 | 0.758 | -0.876 | 0.381 | -2.151 | 0.822 |
| C(Test_Drug)[T.Veh] | 0.0246 | 0.886 | 0.028 | 0.978 | -1.712 | 1.761 |
| C(Pre_Treatment)[T.Sal] : C(Test_Drug)[T.Veh] | 0.2153 | 1.287 | 0.167 | 0.867 | -2.307 | 2.738 |
| Basal_centered | -2.8300 | 1.233 | -2.296 | 0.022 | -5.246 | -0.414 |
| Basal_centered : C(Pre_Treatment)[T.Sal] | 3.1589 | 1.253 | 2.520 | 0.012 | 0.702 | 5.616 |
| Basal_centered : C(Test_Drug)[T.Veh] | 3.4277 | 1.243 | 2.757 | 0.006 | 0.991 | 5.865 |

| | | | | | | |
|---|---|---|---|---|---|---|
| Basal_centered : C(Pre_Treatment)[T.Sal] : C(Test_Drug)[T.Veh] | -3.3264 | 1.352 | -2.461 | 0.014 | -5.976 | -0.677 |

**Table 3**. Statistical analysis of Fig. 3A.

| 3x2 Standard ANOVA (Pingouin, Type III SS) | | |
|---|---|---|
| Effect | F (DFn, DFd) | p |
| Pre_Treatment | F(2,32) = 10.4675 | 0.0003 |
| Test_Drug | F(1,32) = 0.0758 | 0.7848 |
| Pre_Treatment * Test_Drug | F(2,32) = 0.4683 | 0.6303 |

**Table 4**. Statistical analysis of Fig. 3C.

| Linear Mixed Model (Days 1,5,7 Sens vs Non) | |
|---|---|
| Metric | Value |
| Model | MixedLM |
| Dependent Variable | Locomotor |
| No. Observations | 57 |
| No. Groups | 19 |
| Method | REML |
| Scale | 366.5422 |
| Min. group size | 3 |
| Max. group size | 3 |
| Mean group size | 3.0 |
| Log-Likelihood | -247.6037 |
| Converged | Yes |

| Effect | Coef. | Std. Err. | z | p | 95% CI (Lower) | 95% CI (Upper) |
|---|---|---|---|---|---|---|
| Intercept | 89.662 | 19.745 | 4.541 | 0.000 | 50.962 | 128.361 |
| C(Status)[T.Sensitized] | -36.146 | 29.618 | -1.220 | 0.222 | -94.196 | 21.903 |
| C(Test_Drug)[T.Veh] | -10.784 | 26.735 | -0.403 | 0.687 | -63.183 | 41.616 |
| C(Status)[T.Sensitized]:C(Test_Drug)[T.Veh] | 2.786 | 41.103 | 0.068 | 0.946 | -77.774 | 83.345 |
| Day | -5.302 | 1.982 | -2.676 | 0.007 | -9.186 | -1.418 |
| Day:C(Status)[T.Sensitized] | 13.973 | 2.973 | 4.700 | 0.000 | 8.146 | 19.799 |
| Day:C(Test_Drug)[T.Veh] | 1.208 | 2.683 | 0.450 | 0.653 | -4.051 | 6.467 |
| Day:C(Status)[T.Sensitized]:C(Test_Drug)[T.Veh] | -4.750 | 4.125 | -1.152 | 0.250 | -12.836 | 3.335 |

**Table 5**. Statistical analysis of Fig. 3D.

| 2x2 Standard ANOVA (Pingouin, Type III SS) | | |
|---|---|---|
| Effect | F (DFn, DFd) | p |

| | | |
|---|---|---|
| Treatment | F(1,15) = 3.5067 | 0.0807 |
| Drug | F(1,15) = 0.0257 | 0.8747 |
| Interaction | F(1,15) = 0.6263 | 0.4411 |

**Table 6**. Statistical analysis of Fig. 4A.

| D2R in DLS (Global Factorial LMM) | |
|---|---|
| Metric | Value |
| Model | MixedLM |
| Dependent Variable | Raw_Protein_Scaled |
| No. Observations | 113 |
| No. Groups | 11 |
| Method | REML |
| Scale | 0.0562 |
| Min. group size | 7 |
| Max. group size | 12 |
| Mean group size | 10.3 |
| Log-Likelihood | -15.3624 |
| Converged | Yes |

| Effect | Coef. | Std. Err. | z | p | 95% CI (Lower) | 95% CI (Upper) |
|---|---|---|---|---|---|---|
| Intercept | 0.063 | 0.186 | 0.340 | 0.734 | -0.301 | 0.427 |
| C(Group, Sum)[S.AMPH_A] | -0.050 | 0.037 | -1.355 | 0.175 | -0.122 | 0.022 |
| C(Group, Sum)[S.AMPH_R] | 0.032 | 0.034 | 0.930 | 0.353 | -0.035 | 0.099 |
| C(Injection, Sum)[S.Cmpd] | -0.005 | 0.025 | -0.189 | 0.850 | -0.054 | 0.044 |
| C(Group, Sum)[S.AMPH_A]:C(Injection, Sum)[S.Cmpd] | 0.057 | 0.034 | 1.705 | 0.088 | -0.009 | 0.123 |
| C(Group, Sum)[S.AMPH_R]:C(Injection, Sum)[S.Cmpd] | -0.026 | 0.031 | -0.834 | 0.404 | -0.088 | 0.035 |
| Ponceau_Scaled | 0.936 | 0.181 | 5.166 | 0.000 | 0.581 | 1.291 |
| Group Var | 0.009 | 0.029 | | | | |

**Table 7**. Statistical analysis of Fig. 4B.

| D2R in DMS (Global Factorial LMM) | |
|---|---|
| Metric | Value |
| Model | MixedLM |
| Dependent Variable | Raw_Protein_Scaled |
| No. Observations | 119 |
| No. Groups | 12 |
| Method | REML |
| Scale | 0.1138 |

| | |
|---|---|
| Min. group size | 8 |
| Max. group size | 12 |
| Mean group size | 9.9 |
| Log-Likelihood | -55.4483 |
| Converged | Yes |

| Effect | Coef. | Std. Err. | z | p | 95% CI (Lower) | 95% CI (Upper) |
|---|---|---|---|---|---|---|
| Intercept | -0.758 | 0.257 | -2.946 | 0.003 | -1.262 | -0.254 |
| C(Group, Sum)[S.AMPH_A] | 0.116 | 0.051 | 2.254 | 0.024 | 0.015 | 0.216 |
| C(Group, Sum)[S.AMPH_R] | 0.148 | 0.048 | 3.063 | 0.002 | 0.053 | 0.242 |
| C(Injection, Sum)[S.Cmpd] | 0.005 | 0.034 | 0.156 | 0.876 | -0.061 | 0.071 |
| C(Group, Sum)[S.AMPH_A]:C(Injection, Sum)[S.Cmpd] | 0.083 | 0.047 | 1.763 | 0.078 | -0.009 | 0.175 |
| C(Group, Sum)[S.AMPH_R]:C(Injection, Sum)[S.Cmpd] | -0.033 | 0.042 | -0.782 | 0.434 | -0.116 | 0.050 |
| Ponceau_Scaled | 1.698 | 0.255 | 6.663 | 0.000 | 1.199 | 2.197 |
| Group Var | 0.018 | 0.042 | | | | |

**Table 8**. Statistical analysis of Fig. 4C.

| D2R in DS syn (Global Factorial LMM) | |
|---|---|
| Metric | Value |
| Model | MixedLM |
| Dependent Variable | Raw_Protein_Scaled |
| No. Observations | 96 |
| No. Groups | 9 |
| Method | REML |
| Scale | 0.0875 |
| Min. group size | 8 |
| Max. group size | 12 |
| Mean group size | 10.7 |
| Log-Likelihood | -38.5373 |
| Converged | Yes |

| Effect | Coef. | Std. Err. | z | p | 95% CI (Lower) | 95% CI (Upper) |
|---|---|---|---|---|---|---|
| Intercept | 0.205 | 0.238 | 0.863 | 0.388 | -0.261 | 0.672 |
| C(Group, Sum)[S.AMPH_A] | -0.031 | 0.054 | -0.574 | 0.566 | -0.136 | 0.075 |
| C(Group, Sum)[S.AMPH_R] | 0.029 | 0.047 | 0.623 | 0.533 | -0.063 | 0.122 |
| C(Injection, Sum)[S.Cmpd] | -0.028 | 0.032 | -0.883 | 0.377 | -0.091 | 0.035 |

| | | | | | | |
|---|---|---|---|---|---|---|
| C(Group, Sum)[S.AMPH_A]:C(Injection, Sum)[S.Cmpd] | 0.079 | 0.046 | 1.713 | 0.087 | -0.011 | 0.170 |
| C(Group, Sum)[S.AMPH_R]:C(Injection, Sum)[S.Cmpd] | -0.038 | 0.042 | -0.905 | 0.365 | -0.121 | 0.044 |
| Ponceau_Scaled | 0.772 | 0.222 | 3.476 | 0.001 | 0.337 | 1.207 |
| Group Var | 0.056 | 0.118 | | | | |

**Table 9**. Statistical analysis of Fig. 4D.

| Metric | Value |
|---|---|
| Model | MixedLM |
| Dependent Variable | Raw_Protein_Scaled |
| No. Observations | 36 |
| No. Groups | 4 |
| Method | REML |
| Scale | 0.1360 |
| Min. group size | 6 |
| Max. group size | 10 |
| Mean group size | 9.0 |
| Log-Likelihood | -22.8378 |
| Converged | Yes |

| Effect | Coef. | Std. Err. | z | p | 95% CI (Lower) | 95% CI (Upper) |
|---|---|---|---|---|---|---|
| Intercept | 0.396 | 0.328 | 1.207 | 0.228 | -0.247 | 1.039 |
| C(Group, Sum)[S.AMPH_A] | -0.104 | 0.064 | -1.624 | 0.104 | -0.230 | 0.022 |
| C(Injection, Sum)[S.Cmpd] | 0.046 | 0.063 | 0.729 | 0.466 | -0.077 | 0.169 |
| C(Group, Sum)[S.AMPH_A]:C(Injection, Sum)[S.Cmpd] | 0.030 | 0.063 | 0.481 | 0.630 | -0.094 | 0.155 |
| Ponceau_Scaled | 0.577 | 0.300 | 1.926 | 0.054 | -0.010 | 1.164 |
| Group Var | 0.064 | 0.189 | | | | |

**Table 10**. Statistical analysis of Fig. 4E.

| GRK2 in DLS (Global Factorial LMM) | |
|---|---|
| Metric | Value |
| Model | MixedLM |
| Dependent Variable | Raw_Protein_Scaled |
| No. Observations | 169 |
| No. Groups | 16 |
| Method | REML |
| Scale | 0.0488 |

| | |
|---|---|
| Min. group size | 7 |
| Max. group size | 12 |
| Mean group size | 10.6 |
| Log-Likelihood | -6.3089 |
| Converged | No |

| Effect | Coef. | Std. Err. | z | p | 95% CI (Lower) | 95% CI (Upper) |
|---|---|---|---|---|---|---|
| Intercept | 0.459 | 0.110 | 4.188 | 0.000 | 0.244 | 0.674 |
| C(Group, Sum)[S.AMPH_A] | 0.025 | 0.028 | 0.879 | 0.379 | -0.031 | 0.080 |
| C(Group, Sum)[S.AMPH_R] | 0.027 | 0.025 | 1.078 | 0.281 | -0.022 | 0.076 |
| C(Injection, Sum)[S.Cmpd] | -0.006 | 0.018 | -0.363 | 0.717 | -0.041 | 0.028 |
| C(Group, Sum)[S.AMPH_A]:C(Injection, Sum)[S.Cmpd] | -0.002 | 0.026 | -0.086 | 0.932 | -0.053 | 0.048 |
| C(Group, Sum)[S.AMPH_R]:C(Injection, Sum)[S.Cmpd] | 0.033 | 0.023 | 1.406 | 0.160 | -0.013 | 0.078 |
| Ponceau_Scaled | 0.533 | 0.107 | 4.994 | 0.000 | 0.324 | 0.742 |
| Group Var | 0.006 | 0.024 | | | | |

**Table 11**. Statistical analysis of Fig. 4F.

| GRK2 in DMS (Global Factorial LMM) | |
|---|---|
| Metric | Value |
| Model | MixedLM |
| Dependent Variable | Raw_Protein_Scaled |
| No. Observations | 164 |
| No. Groups | 16 |
| Method | REML |
| Scale | 0.0749 |
| Min. group size | 7 |
| Max. group size | 12 |
| Mean group size | 10.2 |
| Log-Likelihood | -51.7111 |
| Converged | Yes |

| Effect | Coef. | Std. Err. | z | p | 95% CI (Lower) | 95% CI (Upper) |
|---|---|---|---|---|---|---|
| Intercept | 0.553 | 0.164 | 3.363 | 0.001 | 0.231 | 0.875 |
| C(Group, Sum)[S.AMPH_A] | 0.070 | 0.037 | 1.873 | 0.061 | -0.003 | 0.143 |
| C(Group, Sum)[S.AMPH_R] | 0.028 | 0.034 | 0.820 | 0.412 | -0.039 | 0.094 |
| C(Injection, Sum)[S.Cmpd] | -0.038 | 0.023 | -1.675 | 0.094 | -0.083 | 0.006 |

| | | | | | | |
|---|---|---|---|---|---|---|
| C(Group, Sum)[S.AMPH_A]:C(Injection, Sum)[S.Cmpd] | 0.041 | 0.033 | 1.249 | 0.212 | -0.023 | 0.105 |
| C(Group, Sum)[S.AMPH_R]:C(Injection, Sum)[S.Cmpd] | 0.079 | 0.030 | 2.652 | 0.008 | 0.021 | 0.137 |
| Ponceau_Scaled | 0.441 | 0.148 | 2.974 | 0.003 | 0.151 | 0.732 |
| Group Var | 0.074 | 0.121 | | | | |

**Table 12**. Statistical analysis of Fig. 4G.

| GRK2 in DS syn (Global Factorial LMM) | |
|---|---|
| Metric | Value |
| Model | MixedLM |
| Dependent Variable | Raw_Protein_Scaled |
| No. Observations | 159 |
| No. Groups | 16 |
| Method | REML |
| Scale | 0.0561 |
| Min. group size | 7 |
| Max. group size | 12 |
| Mean group size | 9.9 |
| Log-Likelihood | -23.4557 |
| Converged | Yes |

| Effect | Coef. | Std. Err. | z | p | 95% CI (Lower) | 95% CI (Upper) |
|---|---|---|---|---|---|---|
| Intercept | 0.313 | 0.127 | 2.472 | 0.013 | 0.065 | 0.562 |
| C(Group, Sum)[S.AMPH_A] | -0.008 | 0.032 | -0.244 | 0.807 | -0.071 | 0.056 |
| C(Group, Sum)[S.AMPH_R] | 0.052 | 0.029 | 1.772 | 0.076 | -0.006 | 0.109 |
| C(Injection, Sum)[S.Cmpd] | 0.010 | 0.020 | 0.501 | 0.616 | -0.028 | 0.048 |
| C(Group, Sum)[S.AMPH_A]:C(Injection, Sum)[S.Cmpd] | 0.011 | 0.028 | 0.402 | 0.687 | -0.044 | 0.067 |
| C(Group, Sum)[S.AMPH_R]:C(Injection, Sum)[S.Cmpd] | -0.008 | 0.026 | -0.319 | 0.750 | -0.059 | 0.043 |
| Ponceau_Scaled | 0.677 | 0.120 | 5.640 | 0.000 | 0.442 | 0.913 |
| Group Var | 0.024 | 0.050 | | | | |

**Table 13**. Statistical analysis of Fig. 4H.

| GRK2 in NAc syn (Global Factorial LMM) | |
|---|---|
| Metric | Value |
| Model | MixedLM |

| | |
|---|---|
| Dependent Variable | Raw_Protein_Scaled |
| No. Observations | 131 |
| No. Groups | 14 |
| Method | REML |
| Scale | 0.0887 |
| Min. group size | 6 |
| Max. group size | 12 |
| Mean group size | 9.4 |
| Log-Likelihood | -47.4273 |
| Converged | Yes |

| Effect | Coef. | Std. Err. | z | p | 95% CI (Lower) | 95% CI (Upper) |
|---|---|---|---|---|---|---|
| Intercept | 0.582 | 0.117 | 4.995 | 0.000 | 0.354 | 0.811 |
| C(Group, Sum)[S.AMPH_A] | 0.016 | 0.046 | 0.342 | 0.732 | -0.074 | 0.106 |
| C(Group, Sum)[S.AMPH_R] | 0.007 | 0.040 | 0.186 | 0.853 | -0.071 | 0.085 |
| C(Injection, Sum)[S.Cmpd] | -0.003 | 0.028 | -0.093 | 0.926 | -0.058 | 0.053 |
| C(Group, Sum)[S.AMPH_A]:C(Injection, Sum)[S.Cmpd] | 0.092 | 0.041 | 2.263 | 0.024 | 0.012 | 0.172 |
| C(Group, Sum)[S.AMPH_R]:C(Injection, Sum)[S.Cmpd] | -0.116 | 0.036 | -3.233 | 0.001 | -0.187 | -0.046 |
| Ponceau_Scaled | 0.409 | 0.105 | 3.906 | 0.000 | 0.204 | 0.614 |
| Group Var | 0.021 | 0.046 | | | | |

**Table 14**. Statistical analysis of Fig. 4I.

| GRK5 in DLS (Global Factorial LMM) | |
|---|---|
| Metric | Value |
| Model | MixedLM |
| Dependent Variable | Raw_Protein_Scaled |
| No. Observations | 161 |
| No. Groups | 16 |
| Method | REML |
| Scale | 0.0605 |
| Min. group size | 8 |
| Max. group size | 12 |
| Mean group size | 10.1 |
| Log-Likelihood | -22.9564 |
| Converged | No |

| Effect | Coef. | Std. Err. | z | p | 95% CI (Lower) | 95% CI (Upper) |
|---|---|---|---|---|---|---|
| Intercept | 0.191 | 0.126 | 1.512 | 0.130 | -0.057 | 0.438 |

| | | | | | | |
|---|---|---|---|---|---|---|
| C(Group, Sum)[S.AMPH_A] | 0.043 | 0.032 | 1.359 | 0.174 | -0.019 | 0.106 |
| C(Group, Sum)[S.AMPH_R] | 0.070 | 0.028 | 2.506 | 0.012 | 0.015 | 0.125 |
| C(Injection, Sum)[S.Cmpd] | -0.027 | 0.020 | -1.327 | 0.184 | -0.067 | 0.013 |
| C(Group, Sum)[S.AMPH_A]:C(Injection, Sum)[S.Cmpd] | 0.044 | 0.030 | 1.496 | 0.135 | -0.014 | 0.103 |
| C(Group, Sum)[S.AMPH_R]:C(Injection, Sum)[S.Cmpd] | 0.031 | 0.027 | 1.180 | 0.238 | -0.021 | 0.084 |
| Ponceau_Scaled | 0.786 | 0.123 | 6.414 | 0.000 | 0.546 | 1.027 |
| Group Var | 0.007 | 0.019 | | | | |

**Table 15**. Statistical analysis of Fig. 4J.

| GRK5 in DMS (Global Factorial LMM) | |
|---|---|
| Metric | Value |
| Model | MixedLM |
| Dependent Variable | Raw_Protein_Scaled |
| No. Observations | 168 |
| No. Groups | 16 |
| Method | REML |
| Scale | 0.0758 |
| Min. group size | 8 |
| Max. group size | 12 |
| Mean group size | 10.5 |
| Log-Likelihood | -53.5166 |
| Converged | Yes |

| Effect | Coef. | Std. Err. | z | p | 95% CI (Lower) | 95% CI (Upper) |
|---|---|---|---|---|---|---|
| Intercept | 0.428 | 0.167 | 2.568 | 0.010 | 0.101 | 0.755 |
| C(Group, Sum)[S.AMPH_A] | 0.141 | 0.037 | 3.815 | 0.000 | 0.068 | 0.213 |
| C(Group, Sum)[S.AMPH_R] | 0.119 | 0.034 | 3.513 | 0.000 | 0.053 | 0.185 |
| C(Injection, Sum)[S.Cmpd] | -0.009 | 0.022 | -0.417 | 0.676 | -0.053 | 0.035 |
| C(Group, Sum)[S.AMPH_A]:C(Injection, Sum)[S.Cmpd] | 0.040 | 0.032 | 1.242 | 0.214 | -0.023 | 0.103 |
| C(Group, Sum)[S.AMPH_R]:C(Injection, Sum)[S.Cmpd] | -0.008 | 0.029 | -0.279 | 0.781 | -0.066 | 0.050 |
| Ponceau_Scaled | 0.551 | 0.150 | 3.663 | 0.000 | 0.256 | 0.846 |
| Group Var | 0.076 | 0.118 | | | | |

**Table 16**. Statistical analysis of Fig. 4K.

| GRK5 in DS syn (Global Factorial LMM) | |
|---|---|
| Metric | Value |
| Model | MixedLM |
| Dependent Variable | Raw_Protein_Scaled |
| No. Observations | 157 |
| No. Groups | 16 |
| Method | REML |
| Scale | 0.0646 |
| Min. group size | 6 |
| Max. group size | 11 |
| Mean group size | 9.8 |
| Log-Likelihood | -35.6904 |
| Converged | Yes |

| Effect | Coef. | Std. Err. | z | p | 95% CI (Lower) | 95% CI (Upper) |
|---|---|---|---|---|---|---|
| Intercept | 0.386 | 0.146 | 2.637 | 0.008 | 0.099 | 0.673 |
| C(Group, Sum)[S.AMPH_A] | 0.033 | 0.034 | 0.981 | 0.326 | -0.033 | 0.100 |
| C(Group, Sum)[S.AMPH_R] | 0.065 | 0.034 | 1.941 | 0.052 | -0.001 | 0.131 |
| C(Injection, Sum)[S.Cmpd] | 0.008 | 0.021 | 0.380 | 0.704 | -0.033 | 0.049 |
| C(Group, Sum)[S.AMPH_A]:C(Injection, Sum)[S.Cmpd] | 0.037 | 0.030 | 1.251 | 0.211 | -0.021 | 0.096 |
| C(Group, Sum)[S.AMPH_R]:C(Injection, Sum)[S.Cmpd] | -0.031 | 0.029 | -1.074 | 0.283 | -0.088 | 0.026 |
| Ponceau_Scaled | 0.610 | 0.138 | 4.423 | 0.000 | 0.340 | 0.880 |
| Group Var | 0.036 | 0.067 | | | | |

**Table 17**. Statistical analysis of Fig. 4L.

| GRK5 in NAc syn (Global Factorial LMM) | |
|---|---|
| Metric | Value |
| Model | MixedLM |
| Dependent Variable | Raw_Protein_Scaled |
| No. Observations | 137 |
| No. Groups | 14 |
| Method | REML |
| Scale | 0.1240 |
| Min. group size | 7 |
| Max. group size | 12 |
| Mean group size | 9.8 |
| Log-Likelihood | -68.9707 |
| Converged | No |

| Effect | Coef. | Std. Err. | z | p | 95% CI (Lower) | 95% CI (Upper) |
|---|---|---|---|---|---|---|
| Intercept | 0.333 | 0.125 | 2.657 | 0.008 | 0.087 | 0.579 |
| C(Group, Sum)[S.AMPH_A] | 0.041 | 0.051 | 0.802 | 0.422 | -0.059 | 0.141 |
| C(Group, Sum)[S.AMPH_R] | 0.052 | 0.046 | 1.119 | 0.263 | -0.039 | 0.143 |
| C(Injection, Sum)[S.Cmpd] | 0.038 | 0.033 | 1.146 | 0.252 | -0.027 | 0.102 |
| C(Group, Sum)[S.AMPH_A]:C(Injection, Sum)[S.Cmpd] | 0.155 | 0.046 | 3.375 | 0.001 | 0.065 | 0.245 |
| C(Group, Sum)[S.AMPH_R]:C(Injection, Sum)[S.Cmpd] | -0.114 | 0.042 | -2.712 | 0.007 | -0.197 | -0.032 |
| Ponceau_Scaled | 0.645 | 0.116 | 5.549 | 0.000 | 0.417 | 0.873 |
| Group Var | 0.019 | 0.048 | | | | |

**Table 18**. Statistical analysis of Fig. 5A.

| Metric | Value |
|---|---|
| Model | OLS |
| N | 52 |
| $R^2$ | 0.078 |
| Adj. $R^2$ | 0.018 |
| F-statistic | 1.301 |
| p (F) | 0.282 |
| AIC | 12.44 |
| BIC | 22.96 |
| Covariance | HC3 |

| Variable | Coef | Std.Err | z | p | 95% CI lower | 95% CI upper |
|---|---|---|---|---|---|---|
| Intercept | 0.0725 | 0.192 | 0.378 | 0.705 | -0.304 | 0.449 |
| X_Centered | 0.0983 | 0.214 | 0.459 | 0.646 | -0.321 | 0.518 |
| Is_Cmpd | -0.0062 | 0.066 | -0.094 | 0.925 | -0.136 | 0.123 |
| X_Centered:Is_Cmpd | -0.2157 | 0.254 | -0.850 | 0.395 | -0.714 | 0.283 |
| Ponceau_Scaled | 0.9186 | 0.187 | 4.912 | 0.000 | 0.553 | 1.284 |

**Table 19**. Statistical analysis of Fig. 5B.

| Metric | Value |
|---|---|
| Model | OLS |
| N | 58 |
| $R^2$ | 0.210 |
| Adj. $R^2$ | 0.152 |
| F-statistic | 3.640 |
| p (F) | 0.010 |
| AIC | 45.38 |

| | |
|---|---|
| BIC | 57.02 |
| Covariance | HC3 |

| Variable | Coef | Std.Err | z | p | 95% CI lower | 95% CI upper |
|---|---|---|---|---|---|---|
| Intercept | -0.7421 | 0.261 | -2.842 | 0.004 | -1.254 | -0.230 |
| X_Centered | 0.2213 | 0.214 | 1.033 | 0.302 | -0.198 | 0.641 |
| Is_Cmpd | 0.0061 | 0.074 | 0.082 | 0.935 | -0.139 | 0.151 |
| X_Centered:Is_Cmpd | -0.2874 | 0.268 | -1.073 | 0.283 | -0.813 | 0.238 |
| Ponceau_Scaled | 1.6825 | 0.262 | 6.420 | 0.000 | 1.169 | 2.196 |

**Table 20**. Statistical analysis of Fig. 5C.

| Metric | Value |
|---|---|
| Model | OLS |
| N | 52 |
| $R^2$ | 0.102 |
| Adj. $R^2$ | 0.031 |
| F-statistic | 1.437 |
| p (F) | 0.237 |
| AIC | 28.77 |
| BIC | 40.21 |
| Covariance | HC3 |

| Variable | Coef | Std.Err | z | p | 95% CI lower | 95% CI upper |
|---|---|---|---|---|---|---|
| Intercept | 0.1984 | 0.245 | 0.810 | 0.418 | -0.282 | 0.679 |
| X_Centered | 0.1762 | 0.234 | 0.753 | 0.451 | -0.282 | 0.634 |
| Is_Cmpd | -0.0295 | 0.081 | -0.364 | 0.716 | -0.188 | 0.129 |
| X_Centered:Is_Cmpd | -0.3017 | 0.291 | -1.037 | 0.300 | -0.872 | 0.269 |
| Ponceau_Scaled | 0.7689 | 0.229 | 3.357 | 0.001 | 0.320 | 1.218 |

**Table 21**. Statistical analysis of Fig. 5D.

| Metric | Value |
|---|---|
| Model | OLS |
| N | 36 |
| $R^2$ | 0.118 |
| Adj. $R^2$ | 0.016 |
| F-statistic | 1.157 |
| p (F) | 0.347 |
| AIC | 32.81 |
| BIC | 41.56 |
| Covariance | HC3 |

| Variable | Coef | Std.Err | z | p | 95% CI lower | 95% CI upper |
|---|---|---|---|---|---|---|

| | | | | | | |
|---|---|---|---|---|---|---|
| Intercept | 0.3921 | 0.338 | 1.160 | 0.246 | -0.271 | 1.055 |
| X_Centered | 0.1587 | 0.289 | 0.549 | 0.583 | -0.408 | 0.726 |
| Is_Cmpd | 0.0442 | 0.089 | 0.497 | 0.619 | -0.131 | 0.219 |
| X_Centered:Is_Cmpd | -0.2675 | 0.321 | -0.834 | 0.404 | -0.896 | 0.361 |
| Ponceau_Scaled | 0.5698 | 0.296 | 1.926 | 0.054 | -0.011 | 1.150 |

**Table 22**. Statistical analysis of Fig. 5E.

| Metric | Value |
|---|---|
| Model | OLS |
| N | 71 |
| $R^2$ | 0.163 |
| Adj. $R^2$ | 0.112 |
| F-statistic | 5.412 |
| p (F) | 0.000787 |
| AIC | 3.806 |
| BIC | 15.12 |
| Covariance | HC3 |

| Variable | Coef | Std.Err | z | p | 95% CI lower | 95% CI upper |
|---|---|---|---|---|---|---|
| Intercept | 0.5792 | 0.171 | 3.390 | 0.001 | 0.244 | 0.914 |
| X_Centered | 0.1306 | 0.180 | 0.726 | 0.468 | -0.222 | 0.483 |
| Is_Cmpd | 0.0639 | 0.059 | 1.083 | 0.279 | -0.052 | 0.180 |
| X_Centered:Is_Cmpd | -0.2599 | 0.218 | -1.193 | 0.233 | -0.687 | 0.167 |
| Ponceau_Scaled | 0.4204 | 0.169 | 2.492 | 0.013 | 0.090 | 0.751 |

**Table 23**. Statistical analysis of Fig. 5F.

| Metric | Value |
|---|---|
| Model | OLS |
| N | 74 |
| $R^2$ | 0.089 |
| Adj. $R^2$ | 0.035 |
| F-statistic | 1.642 |
| p (F) | 0.170 |
| AIC | 25.52 |
| BIC | 37.06 |
| Covariance | HC3 |

| Variable | Coef | Std.Err | z | p | 95% CI lower | 95% CI upper |
|---|---|---|---|---|---|---|
| Intercept | 0.5043 | 0.181 | 2.788 | 0.005 | 0.150 | 0.859 |
| X_Centered | 0.0327 | 0.202 | 0.162 | 0.872 | -0.364 | 0.429 |
| Is_Cmpd | -0.0306 | 0.063 | -0.487 | 0.626 | -0.154 | 0.093 |
| X_Centered:Is_Cmpd | -0.1722 | 0.246 | -0.699 | 0.485 | -0.654 | 0.309 |

| | | | | | | |
|---|---|---|---|---|---|---|
| Ponceau_Scaled | 0.5121 | 0.175 | 2.928 | 0.003 | 0.169 | 0.855 |

**Table 24**. Statistical analysis of Fig. 5G.

| Metric | Value |
|---|---|
| Model | OLS |
| N | 73 |
| $R^2$ | 0.060 |
| Adj. $R^2$ | 0.004 |
| F-statistic | 1.073 |
| p (F) | 0.377 |
| AIC | -4.41 |
| BIC | 7.13 |
| Covariance | HC3 |

| Variable | Coef | Std.Err | z | p | 95% CI lower | 95% CI upper |
|---|---|---|---|---|---|---|
| Intercept | 0.2985 | 0.135 | 2.210 | 0.027 | 0.033 | 0.564 |
| X_Centered | 0.1068 | 0.150 | 0.713 | 0.476 | -0.187 | 0.401 |
| Is_Cmpd | 0.0124 | 0.048 | 0.258 | 0.796 | -0.082 | 0.107 |
| X_Centered:Is_Cmpd | -0.1075 | 0.183 | -0.588 | 0.557 | -0.466 | 0.251 |
| Ponceau_Scaled | 0.6543 | 0.140 | 4.670 | 0.000 | 0.380 | 0.928 |

**Table 25**. Statistical analysis of Fig. 5H.

| Metric | Value |
|---|---|
| Model | OLS |
| N | 60 |
| $R^2$ | 0.258 |
| Adj. $R^2$ | 0.204 |
| F-statistic | 4.746 |
| p (F) | 0.002 |
| AIC | 18.41 |
| BIC | 30.06 |
| Covariance | HC3 |

| Variable | Coef | Std.Err | z | p | 95% CI lower | 95% CI upper |
|---|---|---|---|---|---|---|
| Intercept | 0.6112 | 0.146 | 4.193 | 0.000 | 0.326 | 0.896 |
| X_Centered | -0.0215 | 0.183 | -0.118 | 0.906 | -0.380 | 0.337 |
| Is_Cmpd | -0.0108 | 0.060 | -0.181 | 0.856 | -0.128 | 0.106 |
| X_Centered:Is_Cmpd | -0.3956 | 0.215 | -1.839 | 0.066 | -0.817 | 0.026 |
| Ponceau_Scaled | 0.4032 | 0.137 | 2.945 | 0.003 | 0.134 | 0.672 |

**Table 26**. Statistical analysis of Fig. 5I.

| Metric | Value |
|---|---|
| Model | OLS |
| N | 72 |
| $R^2$ | 0.115 |
| Adj. $R^2$ | 0.061 |
| F-statistic | 2.126 |
| p (F) | 0.088 |
| AIC | -0.92 |
| BIC | 10.66 |
| Covariance | HC3 |

| Variable | Coef | Std.Err | z | p | 95% CI lower | 95% CI upper |
|---|---|---|---|---|---|---|
| Intercept | 0.1847 | 0.139 | 1.326 | 0.185 | -0.089 | 0.458 |
| X_Centered | 0.1873 | 0.154 | 1.216 | 0.224 | -0.114 | 0.489 |
| Is_Cmpd | -0.0289 | 0.050 | -0.578 | 0.563 | -0.127 | 0.069 |
| X_Centered:Is_Cmpd | -0.1534 | 0.186 | -0.825 | 0.409 | -0.518 | 0.211 |
| Ponceau_Scaled | 0.7812 | 0.138 | 5.655 | 0.000 | 0.511 | 1.052 |

**Table 27**. Statistical analysis of Fig. 5J.

| Metric | Value |
|---|---|
| Model | OLS |
| N | 75 |
| $R^2$ | 0.231 |
| Adj. $R^2$ | 0.183 |
| F-statistic | 4.835 |
| p (F) | 0.002 |
| AIC | 21.67 |
| BIC | 33.27 |
| Covariance | HC3 |

| Variable | Coef | Std.Err | z | p | 95% CI lower | 95% CI upper |
|---|---|---|---|---|---|---|
| Intercept | 0.4176 | 0.182 | 2.296 | 0.022 | 0.061 | 0.774 |
| X_Centered | 0.2154 | 0.205 | 1.051 | 0.293 | -0.186 | 0.617 |
| Is_Cmpd | -0.0087 | 0.063 | -0.138 | 0.890 | -0.133 | 0.115 |
| X_Centered:Is_Cmpd | -0.2481 | 0.249 | -0.996 | 0.319 | -0.737 | 0.241 |
| Ponceau_Scaled | 0.5629 | 0.179 | 3.140 | 0.002 | 0.211 | 0.915 |

**Table 28**. Statistical analysis of Fig. 5K.

| Metric | Value |
|---|---|
| Model | OLS |
| N | 73 |
| $R^2$ | 0.121 |

| | |
|---|---|
| Adj. $R^2$ | 0.067 |
| F-statistic | 2.240 |
| p (F) | 0.073 |
| AIC | 6.78 |
| BIC | 18.33 |
| Covariance | HC3 |

| Variable | Coef | Std.Err | z | p | 95% CI lower | 95% CI upper |
|---|---|---|---|---|---|---|
| Intercept | 0.3758 | 0.150 | 2.505 | 0.012 | 0.082 | 0.670 |
| X_Centered | 0.1492 | 0.166 | 0.899 | 0.369 | -0.176 | 0.474 |
| Is_Cmpd | 0.0076 | 0.053 | 0.143 | 0.886 | -0.097 | 0.112 |
| X_Centered:Is_Cmpd | -0.1915 | 0.201 | -0.952 | 0.341 | -0.586 | 0.203 |
| Ponceau_Scaled | 0.6087 | 0.146 | 4.170 | 0.000 | 0.323 | 0.895 |

**Table 29**. Statistical analysis of Fig. 5L.

| Metric | Value |
|---|---|
| Model | OLS |
| N | 62 |
| $R^2$ | 0.301 |
| Adj. $R^2$ | 0.246 |
| F-statistic | 5.460 |
| p (F) | 0.001 |
| AIC | 34.12 |
| BIC | 46.04 |
| Covariance | HC3 |

| Variable | Coef | Std.Err | z | p | 95% CI lower | 95% CI upper |
|---|---|---|---|---|---|---|
| Intercept | 0.3421 | 0.138 | 2.480 | 0.013 | 0.072 | 0.612 |
| X_Centered | 0.2745 | 0.176 | 1.558 | 0.119 | -0.071 | 0.620 |
| Is_Cmpd | 0.0412 | 0.057 | 0.723 | 0.470 | -0.071 | 0.153 |
| X_Centered:Is_Cmpd | -0.4218 | 0.209 | -2.019 | 0.044 | -0.831 | -0.013 |
| Ponceau_Scaled | 0.6427 | 0.141 | 4.561 | 0.000 | 0.366 | 0.920 |
